# Genomic-Adjusted Radiation Dose from Bulk RNA Sequencing for Personalized Radiotherapy

**DOI:** 10.64898/2026.05.29.728725

**Authors:** Drew T. Bergman, Jill Durkin, Nikhil Joshi, Steven A. Eschrich, Javier F. Torres-Roca, Jacob G. Scott

## Abstract

Radiotherapy is delivered to more than half of all patients with cancer yet is prescribed using uniform physical doses despite well-established interpatient variability in biological response. The genomic-adjusted radiation dose (GARD), derived from the radiosensitivity index (RSI), integrates tumor transcriptomics with radiation dose to estimate patient-specific treatment effect, and has been clinically validated as a predictor of radiotherapy benefit across diverse disease sites, including breast, lung, head and neck, glioma, sarcoma, rectal, and endometrial cancers. However, further clinical validation and deployment has been limited by reliance on microarray-based expression. Here we develop an RNA sequencing–based formulation of RSI (RSI-seq) and show that it preserves the functional properties of the original model across measurement platforms. RSI-seq maintains concordance with microarray RSI, including preservation of patient rank ordering (pooled Spearman *ρ* = 0.86), and, when integrated into GARD, reproduces predicted changes in biological effect under clinically relevant dose perturbations (*R*^2^ ≥ 0.78 for ΔGARD in both directions). This preservation of interventional prediction is robust to expression noise and invariant to normalization strategy, enabling consistent application across RNA-seq pipelines. Application across the TCGA pan-cancer transcriptomic atlas demonstrates scalability across tumor types, with cohort medians agreeing closely with previously published microarray RSI medians (Spearman *ρ* = 0.68, Pearson *r* = 0.85 across 20 matched cohorts). By bringing a clinically validated radiogenomic dose model into the RNA-sequencing era, RSI-seq makes biologically personalized radiotherapy directly accessible, retrospectively in existing RNA-seq cohorts and prospectively in modern clinical sequencing workflows, across the full range of tumor types treated with radiation.

## Background

Radiotherapy is delivered to more than half of all patients with cancer, yet outcomes vary widely across tumors that look identical at the macroscopic level [1, 2]. Much of this variability is intrinsic to the tumor and reflects biological heterogeneity in radiosensitivity [3, 4]. Quantifying that heterogeneity per patient is the foundation of biologically personalized radiation oncology [5, 6].

The Radiosensitivity Index (RSI) is a 10-gene rank-based linear model derived from Affymetrix HG-HU6800 microarray expression in the NCI-60 cell-line panel and validated across multiple clinical cohorts [7–9]. Its dose-response extension, the Genomic-Adjusted Radiation Dose (GARD), couples RSI with the linear-quadratic radiobiological model and a patient-specific dose schedule to yield a continuous, patient-level estimate of biological radiation effect [10–12]. GARD has been validated as a predictor of clinical outcome and of differential benefit from radiotherapy across many tumor sites, including breast [13–15], head and neck [16–18], glioma [19], pancreatic [20], lung [21, 22], sarcoma [23, 24], rectal [25, 26], endometrial [27], and across the pan-cancer setting [28]. Although GARD has thus been validated across many disease sites, its clinical deployment has been limited by reliance on microarray-based expression, creating a disconnect between a clinically validated radiogenomic model and contemporary RNA sequencing–based clinical workflows [29–31].

Here we develop an RNA sequencing–based formulation of RSI and test whether a clinically validated radiogenomic model can be preserved across measurement platforms. We specifically evaluate not only concordance with the original model, but whether the relationship between tumor genomics and radiation treatment effect, captured through GARD, is maintained under clinically relevant dose perturbations.

## Methods

### Cohort and quality control

Matched microarray and RNA-seq expression were assembled for three TCGA tumor types, glioblastoma (GBM), lung squamous cell carcinoma (LUSC), and serous ovarian adenocarcinoma (OV), via the NCI Genomic Data Commons. RNA-seq was processed with the GDC pipeline using STAR alignment and RSEM quantification [32, 33] and used in transcripts-per-million (TPM) form [34]; FPKM and RPKM produced indistinguishable rank orderings (**Supplementary Figure S2A**). Microarray data were RMA-normalized [35]. Samples passed quality control only if they met platform-specific thresholds on signal quality (microarray mean log-intensity above 5; RNA-seq at least 3,000 detected genes, or the within-study 5th percentile if higher), sample-wise Pearson correlation (above 0.40, or the within-study 5th percentile if higher), and PC1–PC2 Mahalanobis distance (within 3 SD) on both platforms (**Supplementary Figure S1**; **Table 1**).

**Table 1.** Cohort composition and quality control. Three TCGA studies (GBM, LUSC, OV) were assessed for both microarray and RNA-seq data quality; samples passed QC only if they met thresholds on both platforms (Methods).

| Study | N samples (matched) | Microarray + RNA-seq QC passed | Downstream cohort |
| --- | --- | --- | --- |
| GBM | 260 | 177 | 177 |
| LUSC | 130 | 107 | 107 |
| OV | 417 | 353 | 353 |
| **Total** | **807** | **637** | **637** |

### RSI calculation and scaling

RSI was computed using the published 10-gene rank-based linear model [8, 9]:

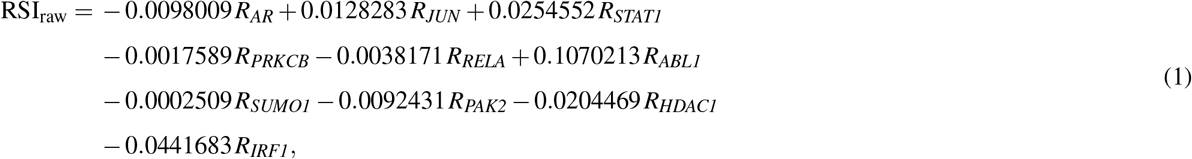

where R_*g*_ is the within-sample rank of gene *g* out of 10 genes. The raw rank-sum has a theoretical achievable range of [−0.605, +1.219] over rank permutations of 1, …, 10 across the 10 RSI genes; we therefore min-max rescale every rank-based RSI value to (0, 1) via

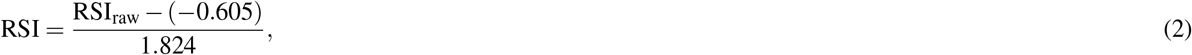

and use the scaled RSI throughout (for both microarray RSI-MA and RNA-seq RSI-TPM, and as the regression target during model development). The scaling is monotonic, so all rank-based and correlation-based comparisons are unchanged; absolute errors are reported on a comparable, calibrated scale.

### RSI-seq candidates and selection

We restricted RNA-seq–native modeling to seven of the original ten RSI genes ({*AR, JUN, STAT1, PRKCB, ABL1, HDAC1, IRF1*}); *RELA, PAK2*, and *SUMO1* were dropped because of consistently low per-gene microarray/RNA-seq Pearson correlation (Results; **Table 2**; **Supplementary Figure S2B**). Samples were partitioned by stratified sampling into a development set (70%, *n* = 447) and a held-out test set (30%, *n* = 190); 5-fold cross-validation within the development set was likewise stratified by tumor type. We considered three candidate model forms that span the natural design choices for translating a rank-based microarray score to RNA-seq: a linear model on log_2_(TPM + 1), which preserves continuous magnitude information and is the natural RNA-seq analogue of the original linear formulation; a within-sample rank model, which is invariant to any monotonic transformation of TPM and therefore theoretically robust to normalization choices but discards magnitude information; and a logistic-squashed linear model on log_2_(TPM + 1), which retains magnitude information while constraining outputs to the (0, 1) interval required by GARD. All three were fit on the same dev/test split and CV folds and on the same seven retained genes:

**Table 2.**
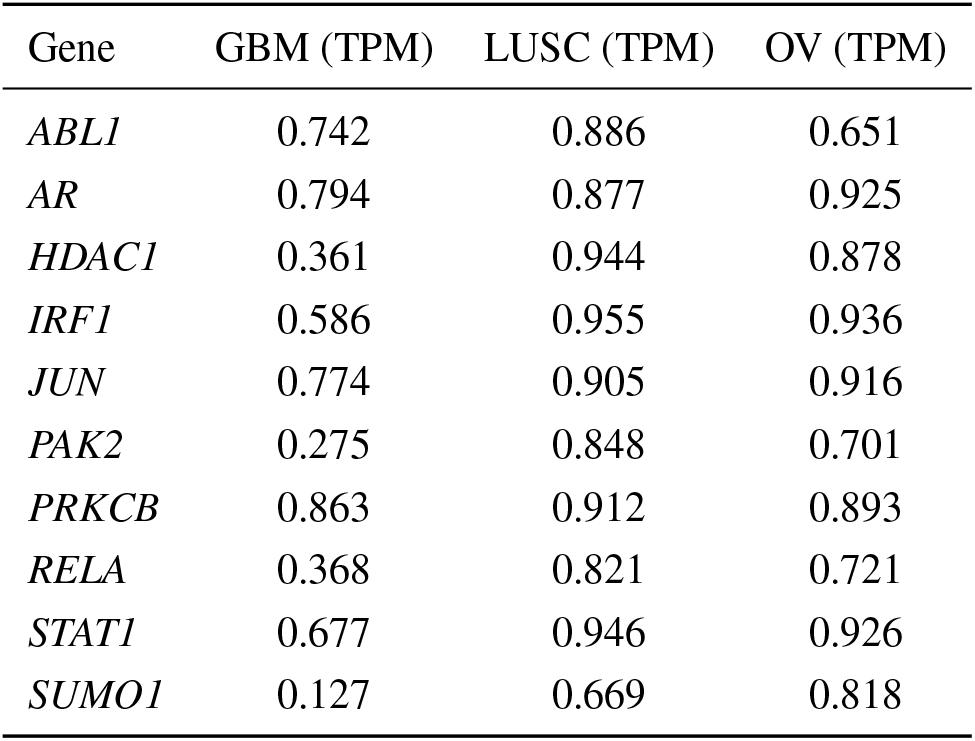
Per-gene Pearson correlations of log_2_-TPM RNA-seq vs. RMA Affymetrix expression across matched biosamples for the ten RSI genes, by TCGA study. *SUMO1, PAK2*, and *RELA* are excluded from the seven-gene RSI-seq because of consistently low cross-platform concordance.

| Gene | GBM (TPM) | LUSC (TPM) | OV (TPM) |
| --- | --- | --- | --- |
| *ABL1* | 0.742 | 0.886 | 0.651 |
| *AR* | 0.794 | 0.877 | 0.925 |
| *HDAC1* | 0.361 | 0.944 | 0.878 |
| *IRF1* | 0.586 | 0.955 | 0.936 |
| *JUN* | 0.774 | 0.905 | 0.916 |
| *PAK2* | 0.275 | 0.848 | 0.701 |
| *PRKCB* | 0.863 | 0.912 | 0.893 |
| *RELA* | 0.368 | 0.821 | 0.721 |
| *STAT1* | 0.677 | 0.946 | 0.926 |
| *SUMO1* | 0.127 | 0.669 | 0.818 |

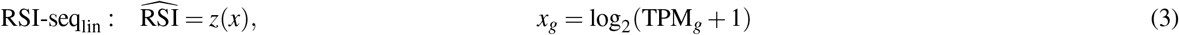

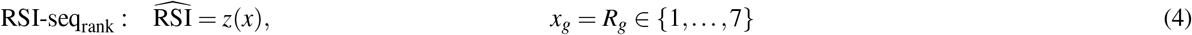

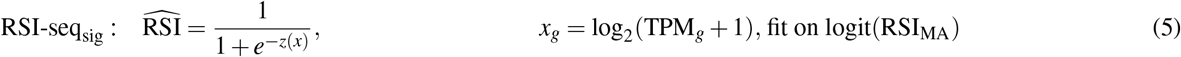

Where 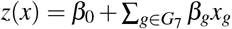 and *G*_7_ = {*AR, JUN, STAT1, PRKCB, ABL1, HDAC1, IRF1*}. For deployability we considered only candidates whose outputs are bounded in (0, 1) by construction, since *α* = −ln(RSI)*/d*_ref_ is undefined as RSI → 0 and goes negative as RSI → 1 and would otherwise destabilize the downstream GARD computation. The unbounded RSI-seq_lin_ is reported alongside its bounded counterparts for transparency but is not deployable in that form. Among the bounded candidates (RSI-seq_rank_ and RSI-seq_sig_), the final model was selected by a composite score combining test MAE, test *R*^2^, GARD MAE, and GARD *R*^2^ with deployability bonuses for invariance to TPM-scale shifts and to monotonic transforms of TPM (**Supplementary Figure S3**); the seven-gene logistic-squashed model (RSI-seq_sig_) was selected, and is referred to as **RSI-seq** hereafter.

### GARD calculation and dose perturbation

For GBM and LUSC samples, in-silico definitive radiotherapy regimens were assigned to match standard-of-care paradigms (60 Gy in 30 fractions). OV was excluded because definitive radiotherapy is not a standard of care in this disease [10]. GARD was computed under the linear-quadratic model:

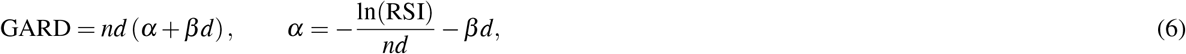

with *β* = 0.05 Gy^−2^ and *d* = 2 Gy. Because GARD is intended to inform dose modulation rather than to fix a single prescription, a practical clinical question is not whether two RSI surrogates yield the same GARD at one fixed dose, but whether they imply the same change in biological effect when dose is modulated, i.e., whether the surrogate is interventionally faithful. To test this, we additionally computed GARD at ±20% of the standard prescription (boost: 72 Gy / 36 fractions; de-escalation: 48 Gy / 24 fractions; both at 2 Gy/fx) and asked, for each sample, whether the per-sample ΔGARD between perturbed and reference regimens implied by each candidate model agrees with the ΔGARD implied by the microarray reference (**Supplementary Figure S6B**; [28]). High concordance on ΔGARD, and not merely on GARD at a single fixed dose, is the property required to use RSI-seq for dose-personalization decisions.

### Pan-TCGA application

To evaluate generalizability beyond the three TCGA cohorts with matched Affymetrix microarray and RNA-seq data, we examined the distribution of RSI-seq across all 32 TCGA pan-cancer atlas cohorts and compared the per-cancer pattern to the previously published pan-cancer microarray RSI analysis derived from the Total Cancer Care (TCC) registry [36]. Per-cancer expression for the seven modeling genes was retrieved through the cBioPortal REST API (the `rna_seq_v2_mrna` molecular profile, RSEM v2 batch-normalized) and passed through **Eq. 5** on a log_2_(*x* + 1) scale. Cohort-level distributions were compared against published per-tumor median microarray RSI from Grass *et al.* [36]; breast subtypes (BC_BASAL, BC_HER2, BC_LUMA, BC_LUMB, BC_NORM) were aggregated into a single BRCA value weighted by reported cohort *N*, COLON+READ_AN were aggregated into CRC, and cohorts without a clean TCGA pan-cancer atlas analogue (LU_NOS, NE, NE_LUNG, NE_PANC, NMSC, KIR_PEL) were excluded.

### Statistical analysis

Differences between dependent overlapping correlations were tested with Steiger’s *Z* [37]; per-sample paired absolute errors with Wilcoxon signed-rank tests; cross-platform and cross-cohort agreement with Spearman *ρ* (rank concordance) and Pearson *r* (linear agreement); calibration with decile means and Bland–Altman limits of agreement (mean ±1.96 SD). Pairwise comparisons across model variants used Bonferroni correction for multiple testing. Robustness to expression noise was evaluated by Gaussian perturbation of log_2_(TPM + 1) (*σ* ∈ {0.10, 0.25, 0.50}, 25 reps per level). Bootstrap 95% confidence intervals on test metrics were obtained from 1,000 paired non-parametric resamples of the test set. Analyses used R version 4.5.1 (2025-06-13) [38].

## Results

### Direct application of the original RSI to RNA-seq introduces a small platform shift

We assembled 807 TCGA biosamples with matched microarray and RNA-seq, and after parallel quality control retained 637 (GBM *n* = 177, LUSC *n* = 107, OV *n* = 353; **Table 1**). Applied directly to RNA-seq TPM, the original 10-gene rank-based RSI agreed reasonably well with microarray RSI (*R*^2^ = 0.65, MAE = 0.087 on the scaled [0, 1] axis) but with a small systematic offset toward lower values (mean shift = 0.064 on the same scale; one-tailed paired Wilcoxon *p <* 0.001; **Figure 1A,B**). The size and consistency of this offset motivated the per-gene cross-platform analysis that follows.

**Figure 1.**
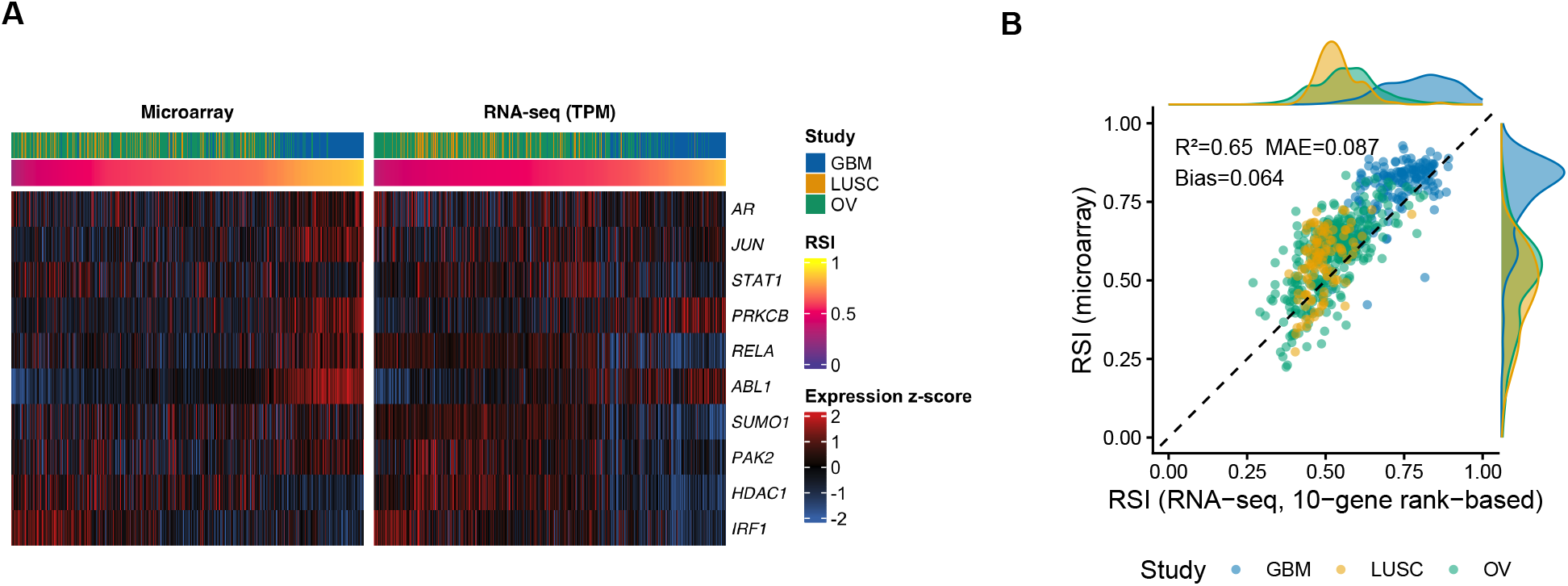
Cross-platform RSI-gene expression and rank-based RSI agreement in matched TCGA biosamples. (**A**) *z*-scored expression of the 10 RSI genes across matched biosamples from TCGA GBM, LUSC, and OV (samples ordered by RSI), rendered side-by-side for Affymetrix microarray (left) and RNA-seq TPM (right). Top annotation bars indicate study and per-sample RSI. (**B**) Scaled RSI computed from rank-based RNA-seq (TPM) vs. Affymetrix microarray (*n* = 637); points colored by study; dashed line is the identity. RSI-TPM agrees well with RSI-MA (*R*^2^ = 0.65, MAE = 0.087) with a small systematic offset toward lower values (mean shift = 0.064; one-tailed paired Wilcoxon *p <* 0.001).

### Cross-platform concordance defines an empirical agreement ceiling

Global cross-platform concordance between microarray and RNA-seq expression profiles was substantial (median Spearman *ρ* ≈ 0.77–0.80 per study across the ~10,000 genes shared between platforms), establishing an empirical upper bound on achievable agreement between measurement platforms. Three RSI genes (*PAK2, RELA, SUMO1*) sit at the low end of this distribution (mean *r* = 0.25–0.55 across studies; **Table 2**), consistent with previously noted probe-level annotation ambiguities and probe-versus sequence-based quantification differences [31]; because the original RSI is rank-based, even a single low-concordance gene perturbs the within-sample ordering and therefore the score, mechanistically explaining the platform offset noted above. Notably, the seven retained RSI genes operate at or above the genome-wide median cross-platform Pearson *r* across cancer types (**Supplementary Figure S2B**), indicating that the RSI signal resides within the most reproducible component of the transcriptome. We restricted all subsequent modeling to the seven retained genes.

### A reduced and reparameterized RSI model recovers cross-platform concordance

Given the quantitative constraints imposed by cross-platform measurement, we derived a reduced and reparameterized RSI model that preserves predictive structure while accommodating RNA sequencing–specific expression characteristics. We evaluated three candidate model forms (**Eqs. 3–5**) chosen to span the natural design choices for a sequencing-native RSI surrogate: a linear model on log_2_(TPM + 1) as the direct RNA-seq analogue of the original linear formulation, retaining continuous magnitude information; a within-sample rank model that mirrors the rank-based encoding of the original RSI and is invariant to any monotonic transformation of TPM (and therefore to choice of normalization), at the cost of magnitude information; and a logistic-squashed linear model on log_2_(TPM + 1), which retains magnitude information while constraining outputs to (0, 1), the bounded interval required for the downstream *α* derivation in GARD. All three were trained on the same 70% development set and evaluated on the same held-out 30% test set. The linear variants substantially improved on the original rank-based RSI applied to TPM (RSI-seq_lin_: test MAE = 0.055, *R*^2^ = 0.78; RSI-seq_sig_: 0.055, *R*^2^ = 0.77); the rank-encoded variant trailed (RSI-seq_rank_: 0.067, *R*^2^ = 0.65) due to magnitude information loss (**Figure 2A**; **Table 3**). Per-cancer test performance was tightly clustered across cancers and prediction bias was small and not systematically tumor-type-specific (**Figure 2B–D**). RSI-seq_lin_ is not deployable because its outputs are not bounded in (0, 1) by construction; we report it for transparency. The bounded RSI-seq (sig) was selected as the final model, with the form

**Table 3.** Held-out test-set performance for the three candidate RSI-seq models on the scaled [0, 1] RSI axis (all on the same seven retained RSI genes; same dev/test split; same 5-fold CV folds). RSI-seq_lin_ produces RSI outside (0, 1) in 25% of test samples and is therefore not deployable for downstream GARD; RSI-seq_sig_ (referred to as RSI-seq throughout) is the final model.

| Variant | Bounded | Test MAE | Test $R^2$ | GARD MAE |
| --- | --- | --- | --- | --- |
| RSI-seq<sub>lin</sub> (not deployable) | no | 0.055 | 0.78 | 2.44 |
| RSI-seq<sub>rank</sub> | yes | 0.067 | 0.65 | 2.83 |
| **RSI-seq (final)** | **yes** | **0.055** | **0.77** | **2.35** |

**Figure 2.**
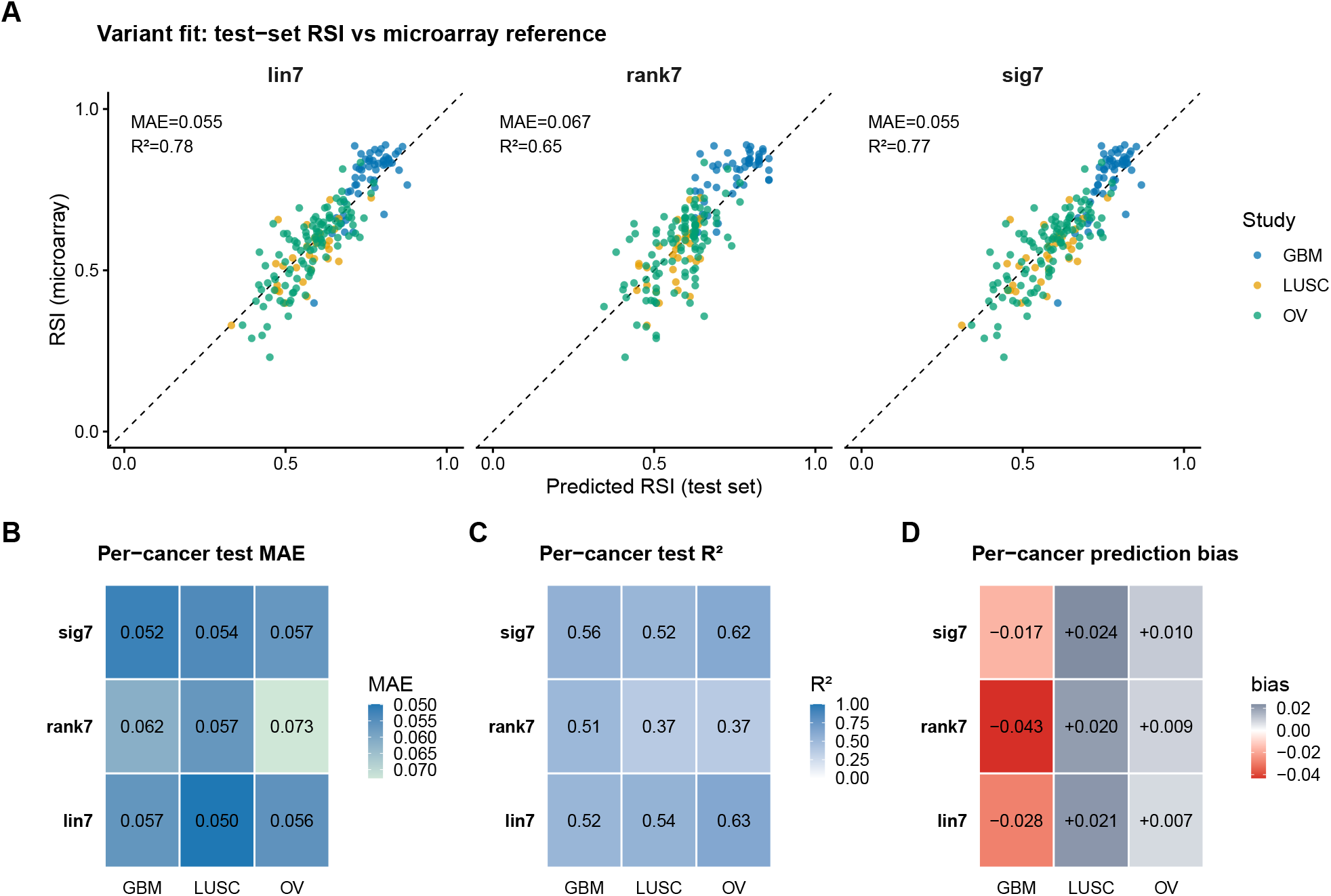
Systematic evaluation of three candidate RSI-seq models. (**A**) Held-out test-set scatter of predicted RSI vs. microarray RSI for each of the three candidate variants (RSI-seq_lin, rank, sig_; *n* = 190). The two TPM-magnitude variants cluster near MAE ≈ 0.055 and *R*^2^ ≈ 0.78; RSI-seq_rank_ trails at MAE 0.067, *R*^2^ = 0.65 due to the information loss of rank encoding. (**B**) Per-cancer test MAE. (**C**) Per-cancer test *R*^2^. (**D**) Per-cancer prediction bias (predicted − microarray RSI). Performance is tightly clustered across cancers, and bias is small and not systematically tumor-type-specific.

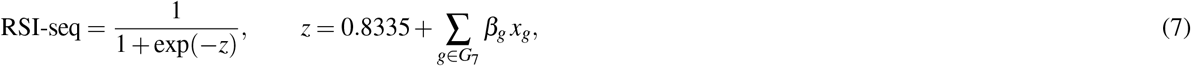

where *x*_*g*_ = log_2_(TPM_*g*_ + 1) and the fitted coefficients are *β*_*AR*_= − 0.0086, *β*_*JUN*_= − 0.0040, *β*_*STAT1*_= + 0.0426, *β*_*PRKCB*_= + 0.0809, *β*_*ABL1*_= + 0.4211, *β*_*HDAC1*_= − 0.2606, *β*_*IRF1*_= − 0.2924. *ABL1, IRF1*, and *HDAC1* remained the dominant contributors with preserved coefficient direction, consistent with the original rank-based RSI [8, 9].

### RSI-seq preserves predicted radiation effect under dose perturbation

To assess whether RSI-seq preserves clinically relevant behavior, we evaluated its impact on predicted radiation effect under dose perturbation. When integrated into GARD under a 60 Gy/30-fx *in silico* regimen, the original RSI applied to RNA-seq (TPM) yielded moderate agreement with GARD-microarray (pooled *R*^2^ = 0.68, MAE = 4.03 in *n* = 284 matched GBM and LUSC samples). RSI-seq substantially improved this to *R*^2^ = 0.78 and MAE = 2.35 (**Figure 3**; **Supplementary Figure S6**), an approximately 42% reduction in GARD MAE relative to the rank-based baseline. Beyond agreement at a single prescription, when dose was perturbed by ±20% (boost: 72 Gy/36 fx; de-escalation: 48 Gy/24 fx; both at 2 Gy/fx), RSI-seq reproduced changes in biological effect under both directions (*R*^2^ ≥ 0.78 for ΔGARD), demonstrating preservation of interventional predictions rather than correlation alone. RSI-seq also preserved the patient ordering implied by RSI-MA across all three TCGA cancers (per-cancer Spearman *ρ* ≥ 0.65, pooled *ρ* = 0.86; **Figure 3A**; **Supplementary Figure S5**).

**Figure 3.**
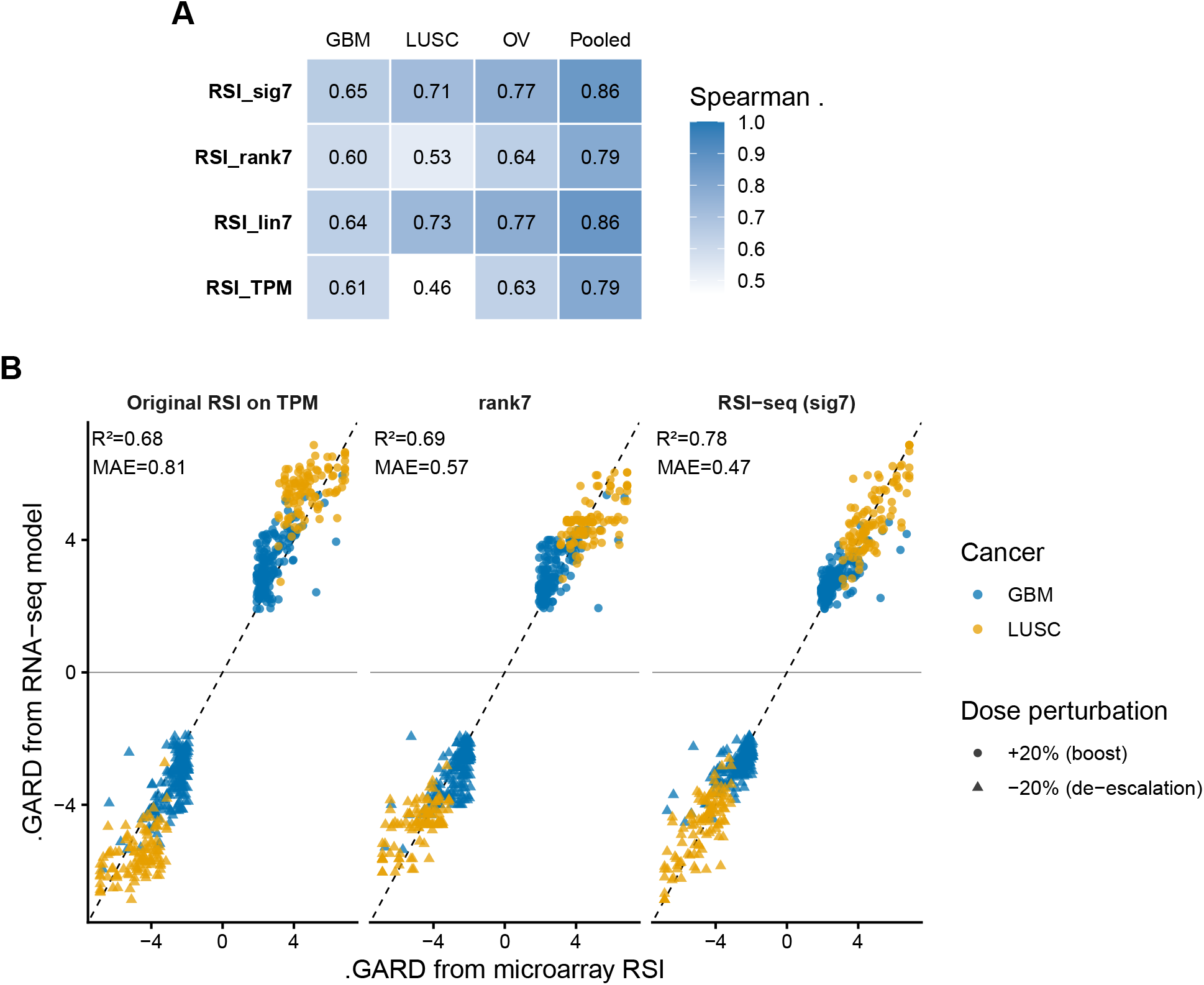
RSI-seq preserves patient ordering and predicted radiation effect under dose perturbation. (**A**) Per-cancer (and pooled) Spearman rank correlation between the patient ordering implied by RSI-MA and the patient ordering implied by each candidate score. The final RSI-seq preserves patient ordering across all three TCGA cancers (per-cancer *ρ* ≥ 0.65; pooled *ρ* = 0.86). (**B**) For each sample, GARD was computed at 60 Gy/30 fx and at ±20% dose modulation (boost: 72 Gy/36 fx; de-escalation: 48 Gy/24 fx; both at 2 Gy/fx). Scatter of ΔGARD predicted by each candidate (y-axis) vs. ΔGARD predicted by RSI-MA (x-axis); circles are boost, triangles are de-escalation. RSI-seq tracks the microarray ΔGARD with *R*^2^ ≥ 0.78 in both directions, substantially better than the original rank-based RSI applied to TPM. Preservation of ΔGARD indicates that RSI-seq maintains the functional relationship between tumor genomics and radiation effect required for treatment decision-making.

### RSI-seq is robust to expression noise and applies across TCGA

We then applied RSI-seq to the RNA-seq data of all 32 TCGA pan-cancer atlas cohorts (n = 10,071 biosamples, cBioPortal, RSEM v2 batch-normalized; Methods) and ordered cohorts by their median RSI-seq (**Figure 4**; **Supplementary Figure S7A**). RSI-seq across cancer spanned the full dynamic range, without any negative values (0.16-0.96). RSI-seq matched expectations based on clinical radiation sensitivity: low-grade glioma (LGG; median 0.89), uveal melanoma (0.88), glioblastoma (0.77), sarcoma (0.76), mesothelioma (0.72), and cutaneous melanoma (0.70) sit at the radioresistant end, while diffuse large B-cell lymphoma (0.44), cervical carcinoma (0.47), and hepatocellular carcinoma (0.52) sit at the radiosensitive end. Per-cancer median RSI-seq agreed with the published per-tumor median microarray RSI of Grass *et al.* [36] (Spearman *ρ* = 0.68, *p* = 8.7 × 10^−4^; Pearson *r* = 0.85, *p* = 2.8 × 10^−6^; *n* = 20 matched cohorts; **Figure 4**). RSI-seq performance was robust to perturbations in gene expression (**Supplementary Figure S4**) and invariant to normalization strategy, with same-sample TPM-vs-RSEM agreement of pooled Spearman *ρ* = 0.97 and Pearson *r* = 0.97 (**Supplementary Figure S7B**). Together, these results indicate that RSI-seq generalizes across tumor types and RNA-seq pipelines and suggest that can be applied directly to any RNA-seq cohort with the seven modeling genes measured.

**Figure 4.**
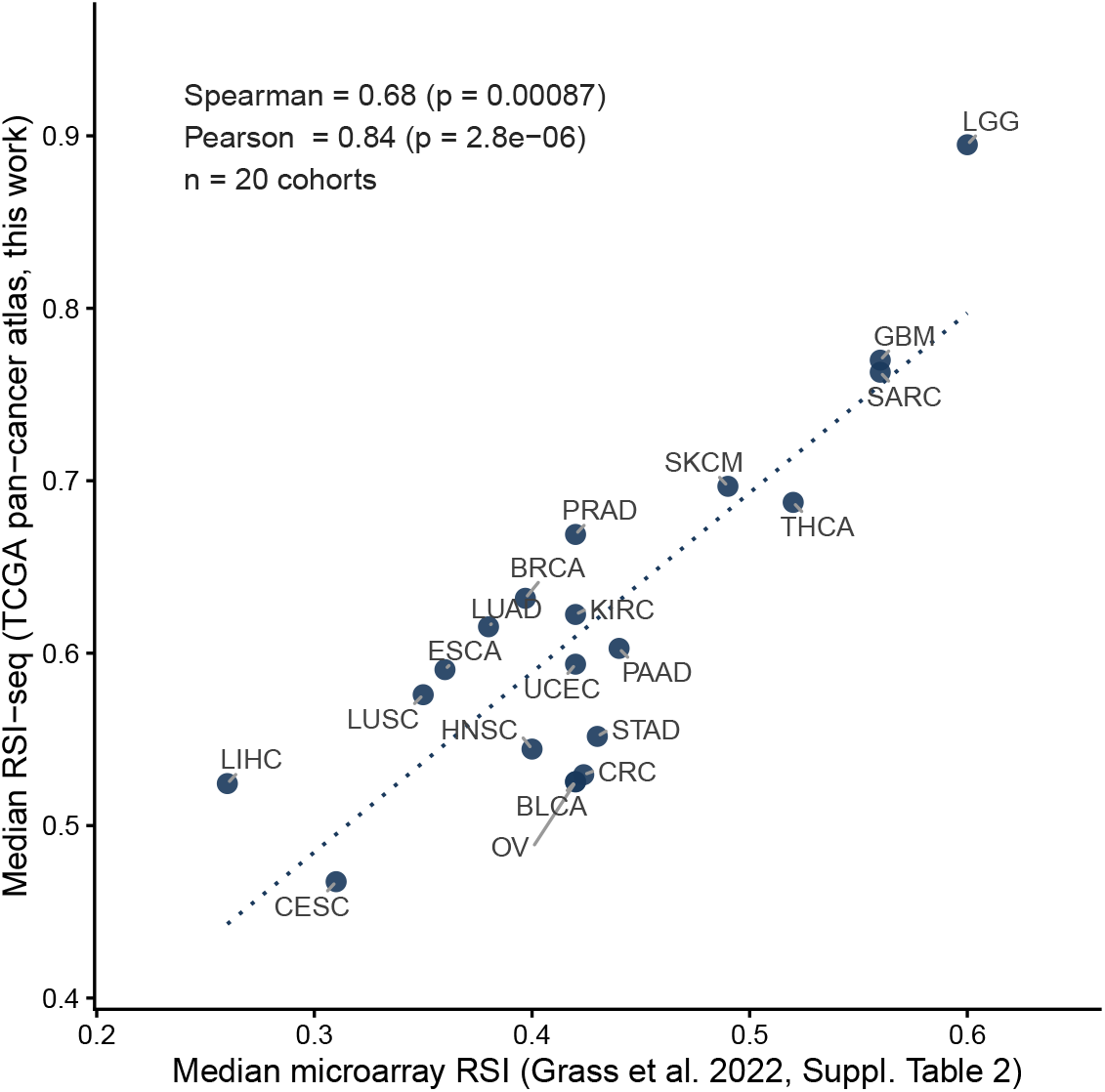
RSI-seq across all 32 TCGA pan-cancer atlas cohorts agrees with published microarray RSI medians. Per-cancer median RSI-seq computed on the cBioPortal pan-cancer atlas RNA-seq (RSEM v2 batch-normalized; COAD and READ merged into CRC; *n* = 20 TCGA cohorts with a clean analogue in the published microarray cohort) plotted against the published per-tumor median microarray RSI [36]. Spearman *ρ* = 0.68 (*p* = 8.7 × 10^−4^), Pearson *r* = 0.85 (*p* = 2.8 × 10^−6^); annotated on the panel. The full per-cohort RSI-seq distribution as a violin plot across all 32 TCGA cancers is shown in Supplementary Figure **S7**A; same-sample TPM-vs-RSEM agreement is shown in Supplementary Figure **S7**B.

## Discussion

These results demonstrate that a clinically validated radiogenomic model can be preserved across measurement platforms without loss of functional behavior. By maintaining the relationship between tumor genomics and radiation treatment effect, RSI-seq enables the deployment of GARD within modern RNA sequencing–based workflows. Compared with the original rank-based RSI applied directly to TPM, RSI-seq lowered test-set MAE from 0.087 to 0.055 (*R*^2^ from 0.65 to 0.77 on the scaled [0, 1] axis), reduced GARD MAE from 4.03 to 2.35 (*R*^2^ from 0.68 to 0.78), preserved patient rank ordering against RSI-MA (per-cancer Spearman *ρ* ≥ 0.65, pooled *ρ* = 0.86), and reproduced the published per-cancer microarray RSI medians across the entire TCGA pan-cancer atlas [36]. Critically, when GARD is computed from RSI-seq, it reproduces the changes in predicted biological effect implied by the microarray reference under clinically realistic ±20% dose perturbations (*R*^2^ ≥ 0.78 in both directions), the property most directly relevant to dose-personalization decisions [10, 28].

The main limitation of this work is the size of the matched-platform development cohort: three TCGA tumor types (GBM, LUSC, OV) provide the parallel microarray and RNA-seq profiling needed to fit and benchmark RSI-seq directly. Although the pan-TCGA application generalizes RSI-seq to all 32 TCGA tumor types and the agreement with published microarray RSI medians is strong, prospective validation against an independent cohort with paired RNA-seq and clinical follow-up — such as the Total Cancer Care registry or other disease-site–specific cohorts that have already been used to validate RSI and GARD on microarray [13–15, 18, 21–27] – remains an important next step.

By removing a key technical barrier to implementation, RSI-seq provides a foundation for prospective evaluation of biologically personalized radiotherapy using routinely acquired transcriptomic data. RSI-seq can be applied to any RNA-seq cohort with the seven modeling genes measured, requires only log_2_(TPM + 1) as input, and returns values in (0, 1) that can be fed directly into the GARD computation. This makes the entire body of clinically validated RSI/GARD evidence [6, 10, 15, 16, 22, 27, 28] immediately accessible to research and clinical programs profiling tumors by RNA sequencing.

## Acknowledgements

The authors thank the TCGA Research Network and the NCI Genomic Data Commons for matched microarray and RNA-seq data. Computational analyses used open-source R and Bioconductor packages.

## Author contributions

DB and JD contributed equally. DB, JD, and JGS designed the study. DB and JD performed data assembly, quality control, and model development. NJ contributed to statistical analysis and model evaluation. SAE and JFTR provided expertise on RSI and GARD; SAE additionally contributed review feedback that motivated the systematic evaluation of candidate models, the genome-wide cross-platform context analysis, the patient sample-rank concordance analysis, and the ΔGARD-under-dose-perturbation analysis. JGS supervised the work. All authors interpreted the results and contributed to the manuscript.

## Data availability

TCGA expression data are publicly available from the NCI Genomic Data Commons (https://gdc.cancer.gov/) and Broad GDAC Firehose (https://gdac.broadinstitute.org/). Code implementing RSI-seq, including the standalone scoring function, fitted coefficients, per-variant performance tables, and the analysis pipeline used to generate the figures, is available from the corresponding author upon reasonable request.

## References

1. Delaney, G., Jacob, S., Featherstone, C. & Barton, M. The role of radiotherapy in cancer treatment: estimating optimal utilization from a review of evidence-based clinical guidelines. Cancer 104, 1129–1137, DOI: 10.1002/cncr.21324 (2005).

2. Baskar, R., Lee, K. A., Yeo, R. & Yeoh, K.-W. Cancer and radiation therapy: current advances and future directions. International Journal Medical Sciences 9, 193–199, DOI: 10.7150/ijms.3635 (2012).

3. Bentzen, S. M. Preventing or reducing late side effects of radiation therapy: radiobiology meets molecular pathology. Nature Reviews Cancer 6, 702–713, DOI: 10.1038/nrc1950 (2006).

4. Hall, E. J. & Giaccia, A. J. Radiobiology for the radiologist. Lippincott Williams & Wilkins, 7th ed. (2012).

5. Poortmans, P., Kaidar-Person, O. & Span, P. Radiation oncology enters the era of individualised medicine. The Lancet Oncology 18, 159–160, DOI: 10.1016/S1470-2045(16)30660-X (2017).

6. Torres-Roca, J. F., Grass, G. D., Scott, J. G. & Eschrich, S. A. Towards data driven rt prescription: Integrating genomics into rt clinical practice. Seminars Radiation Oncology 33, 221–231, DOI: 10.1016/j.semradonc.2023.03.001 (2023).

7. Torres-Roca, J. F. et al. Prediction of radiation sensitivity using a gene expression classifier. Cancer Research 65, 7169–7176, DOI: 10.1158/0008-5472.CAN-05-0656 (2005).

8. Eschrich, S. et al. Systems biology modeling of the radiation sensitivity network: a biomarker discovery platform. International Journal Radiation Oncology*Biology*Physics 75, 497–505, DOI: 10.1016/j.ijrobp.2009.05.056 (2009).

9. Eschrich, S. A. et al. A gene expression model of intrinsic tumor radiosensitivity: prediction of response and prognosis after chemoradiation. International Journal Radiation Oncology*Biology*Physics 75, 489–496, DOI: 10.1016/j.ijrobp.2009.06.014 (2009).

10. Scott, J. G. et al. A genome-based model for adjusting radiotherapy dose (gard): a retrospective, cohort-based study. The Lancet Oncology 18, 202–211, DOI: 10.1016/S1470-2045(16)30648-9 (2017).

11. Fowler, J. F. The linear-quadratic formula and progress in fractionated radiotherapy. British Journal Radiology 62, 679–694, DOI: 10.1259/0007-1285-62-740-679 (1989).

12. Joiner, M. C. & van der Kogel, A. J. Basic clinical radiobiology. CRC Press, 5th ed. (2018).

13. Eschrich, S. A. et al. Validation of a radiosensitivity molecular signature in breast cancer. Clinical Cancer Research 18, 5134–5143, DOI: 10.1158/1078-0432.CCR-12-0891 (2012).

14. Torres-Roca, J. F. et al. Integration of a radiosensitivity molecular signature into the assessment of local recurrence risk in breast cancer. International Journal Radiation Oncology*Biology*Physics 93, 631–638, DOI: 10.1016/j.ijrobp.2015.06.021 (2015).

15. Ahmed, K. A. et al. Utilizing the genomically adjusted radiation dose (gard) to personalize adjuvant radiotherapy in triple negative breast cancer management. EBioMedicine 47, 163–169, DOI: 10.1016/j.ebiom.2019.08.019 (2019).

16. Caudell, J. J. et al. The future of personalised radiotherapy for head and neck cancer. The Lancet Oncology 18, e266–e273, DOI: 10.1016/S1470-2045(17)30252-8 (2017).

17. Cavalieri, S. et al. Clinical validity of a prognostic gene expression cluster-based model in human papillomavirus-positive oropharyngeal carcinoma. JCO Precision Oncology 5, 1666–1676, DOI: 10.1200/PO.21.00094 (2021).

18. Ho, E. et al. Personalized treatment in HPV+ oropharynx cancer using genomic adjusted radiation dose. The Journal Clinical Investigation 135, e194073, DOI: 10.1172/JCI194073 (2025).

19. Ahmed, K. A. et al. Radiosensitivity differences between liver metastases based on primary histology suggest implications for clinical outcomes after stereotactic body radiation therapy. International Journal Radiation Oncology*Biology*Physics 95, 1399–1404, DOI: 10.1016/j.ijrobp.2016.03.050 (2016).

20. Strom, T. et al. Radiosensitivity index predicts for survival with adjuvant radiation in resectable pancreatic cancer. Radiotherapy Oncology 117, 159–164, DOI: 10.1016/j.radonc.2015.07.018 (2015).

21. Ahmed, K. A. et al. Radiosensitivity of lung metastases by primary histology and implications for stereotactic body radiation therapy using the genomically adjusted radiation dose. Journal Thoracic Oncology 13, 1121–1127, DOI: 10.1016/j.jtho.2018.04.027 (2018).

22. Scott, J. G. et al. Personalizing radiotherapy prescription dose using genomic markers of radiosensitivity and normal tissue toxicity in nsclc. Journal Thoracic Oncology 16, 428–438, DOI: 10.1016/j.jtho.2020.11.008 (2021).

23. Yang, G. et al. Genomic identification of sarcoma radiosensitivity and the clinical implications for radiation dose personalization. Translational Oncology 14, 101165, DOI: 10.1016/j.tranon.2021.101165 (2021).

24. Naghavi, A. O. et al. Habitat escalated adaptive therapy (heat): a phase 2 trial utilizing radiomic habitat-directed and genomic-adjusted radiation dose (gard) optimization for high-grade soft tissue sarcoma. BMC Cancer 24, 437, DOI: 10.1186/s12885-024-12151-7 (2024).

25. Yuan, Z. et al. Modeling precision genomic-based radiation dose response in rectal cancer. Future Oncology 16, 2411–2420, DOI: 10.2217/fon-2020-0060 (2020).

26. Xia, H. et al. Validation of a genome-based model for adjusting radiotherapy dose (GARD) in patients with locally advanced rectal cancer. Scientific Reports 14, 21572, DOI: 10.1038/s41598-024-72818-w (2024).

27. Mohammadi, H. et al. Using the radiosensitivity index (rsi) to predict pelvic failure in endometrial cancer treated with adjuvant radiation therapy. International Journal Radiation Oncology*Biology*Physics 106, 496–502, DOI: 10.1016/j.ijrobp.2019.11.013 (2020).

28. Scott, J. G. et al. Pan-cancer prediction of radiotherapy benefit using genomic-adjusted radiation dose (GARD): a cohort-based pooled analysis. The Lancet Oncology 22, 1221–1229, DOI: 10.1016/S1470-2045(21)00347-8 (2021).

29. Stark, R., Grzelak, M. & Hadfield, J. RNA sequencing: the teenage years. Nature Reviews Genetics 20, 631–656, DOI: 10.1038/s41576-019-0150-2 (2019).

30. Marioni, J. C., Mason, C. E., Mane, S. M., Stephens, M. & Gilad, Y. RNA-seq: an assessment of technical reproducibility and comparison with gene expression arrays. Genome Research 18, 1509–1517, DOI: 10.1101/gr.079558.108 (2008).

31. Wang, C. et al. The concordance between RNA-seq and microarray data depends on chemical treatment and transcript abundance. Nature Biotechnology 32, 926–932, DOI: 10.1038/nbt.3001 (2014).

32. Dobin, A. et al. STAR: ultrafast universal RNA-seq aligner. Bioinformatics 29, 15–21, DOI: 10.1093/bioinformatics/bts635 (2013).

33. Li, B. & Dewey, C. N. RSEM: accurate transcript quantification from RNA-seq data with or without a reference genome. BMC Bioinformatics 12, 323, DOI: 10.1186/1471-2105-12-323 (2011).

34. Wagner, G. P., Kin, K. & Lynch, V. J. Measurement of mRNA abundance using RNA-seq data: RPKM measure is inconsistent among samples. Theory Biosciences 131, 281–285, DOI: 10.1007/s12064-012-0162-3 (2012).

35. Irizarry, R. A. et al. Exploration, normalization, and summaries of high density oligonucleotide array probe level data. Biostatistics 4, 249–264, DOI: 10.1093/biostatistics/4.2.249 (2003).

36. Grass, G. D. et al. The radiosensitivity index gene signature identifies distinct tumor immune microenvironment characteristics associated with susceptibility to radiation therapy. International Journal Radiation Oncology*Biology*Physics 113, 635–647, DOI: 10.1016/j.ijrobp.2022.03.006 (2022).

37. Steiger, J. H. Tests for comparing elements of a correlation matrix. Psychological Bulletin 87, 245–251, DOI: 10.1037/0033-2909.87.2.245 (1980).

38. R Core Team. R: A language and environment for statistical computing. R Foundation for Statistical Computing, Vienna, Austria (2024).

