## Supplementary Figures for "Genomic-Adjusted Radiation Dose from Bulk RNA Sequencing for Personalized Radiotherapy"

### Supplementary Information

#### Supplementary Figures

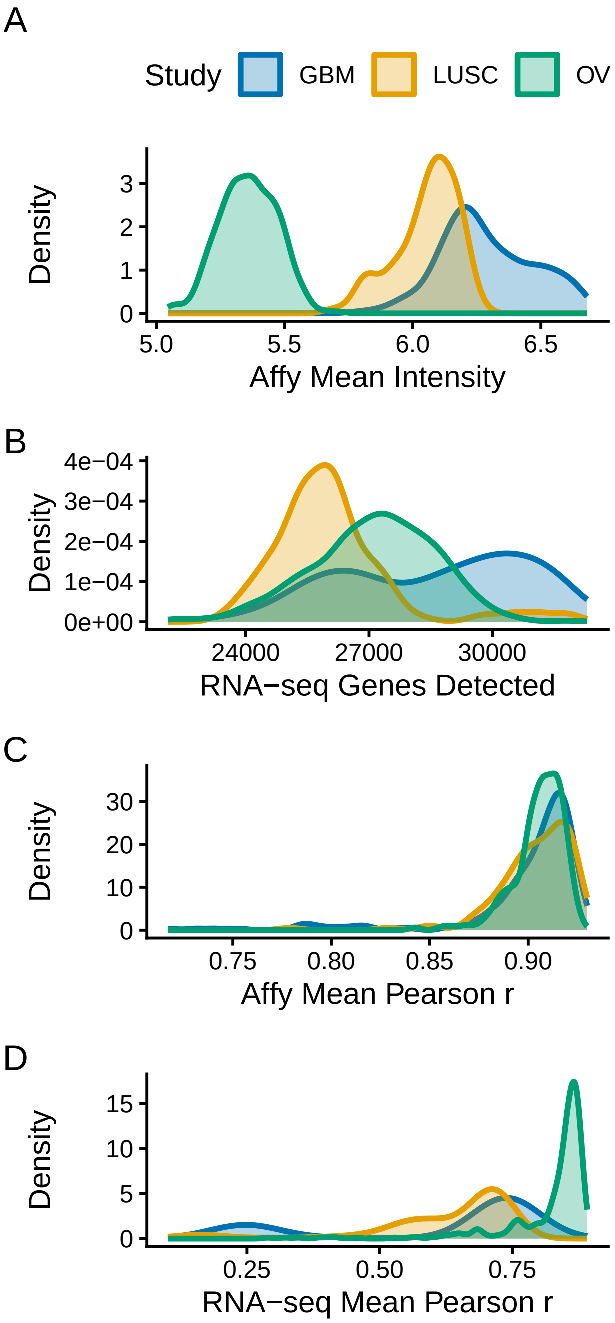


**Figure S1. Per-sample quality-control metric distributions.** Distributions of per-sample quality-control metrics across all TCGA (GBM, LUSC, OV) samples for both Affymetrix microarray and RNA-seq (TPM): mean signal intensity (microarray) or number of detected genes (RNA-seq), mean Pearson correlation with other within-study samples, and Mahalanobis distance in PC1–PC2 space. Inclusion thresholds are described in Methods.

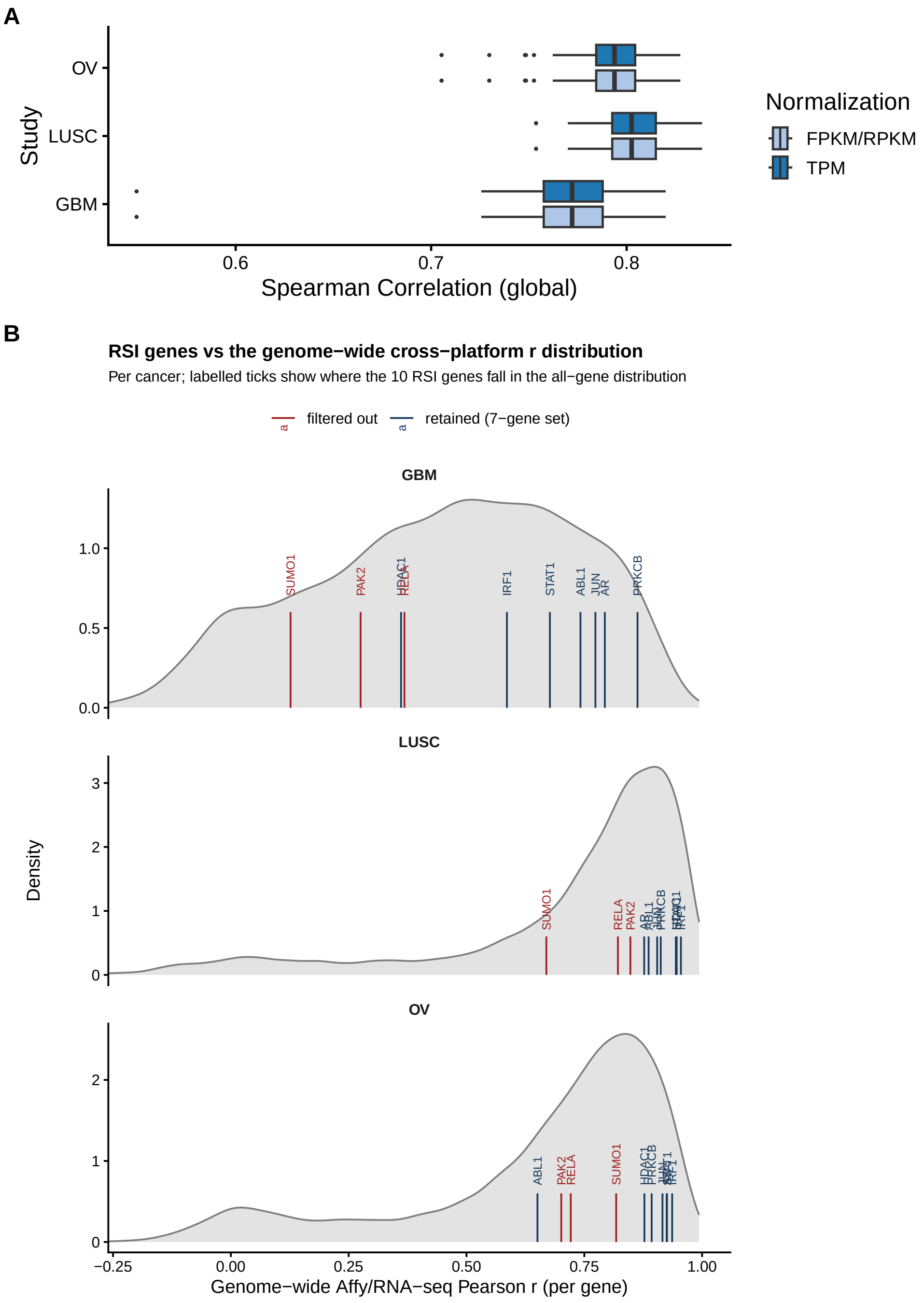


**Figure S2.** **Cross-platform expression characterization.** (**A**) Sample-wise Spearman correlation between matched Affymetrix and RNA-seq (TPM and FPKM/RPKM) profiles across all shared genes per study; Spearman is invariant under monotonic re-normalization, so TPM and FPKM yield identical curves and only TPM is reported throughout. (**B**) For each TCGA cancer, the density of microarray/RNA-seq Pearson *r* across all variance-filtered shared genes (~10,000 genes per cancer; grey). Vertical labelled ticks mark where each of the 10 RSI genes falls in this distribution: the seven retained genes (navy) sit at or above the genome-wide median in every cancer, while the three filtered genes (red; *RELA*, *PAK2*, *SUMO1*) sit at the low end. The retained-gene cross-platform behavior sets an empirical upper bound on the achievable agreement of any RNA-seq RSI model.

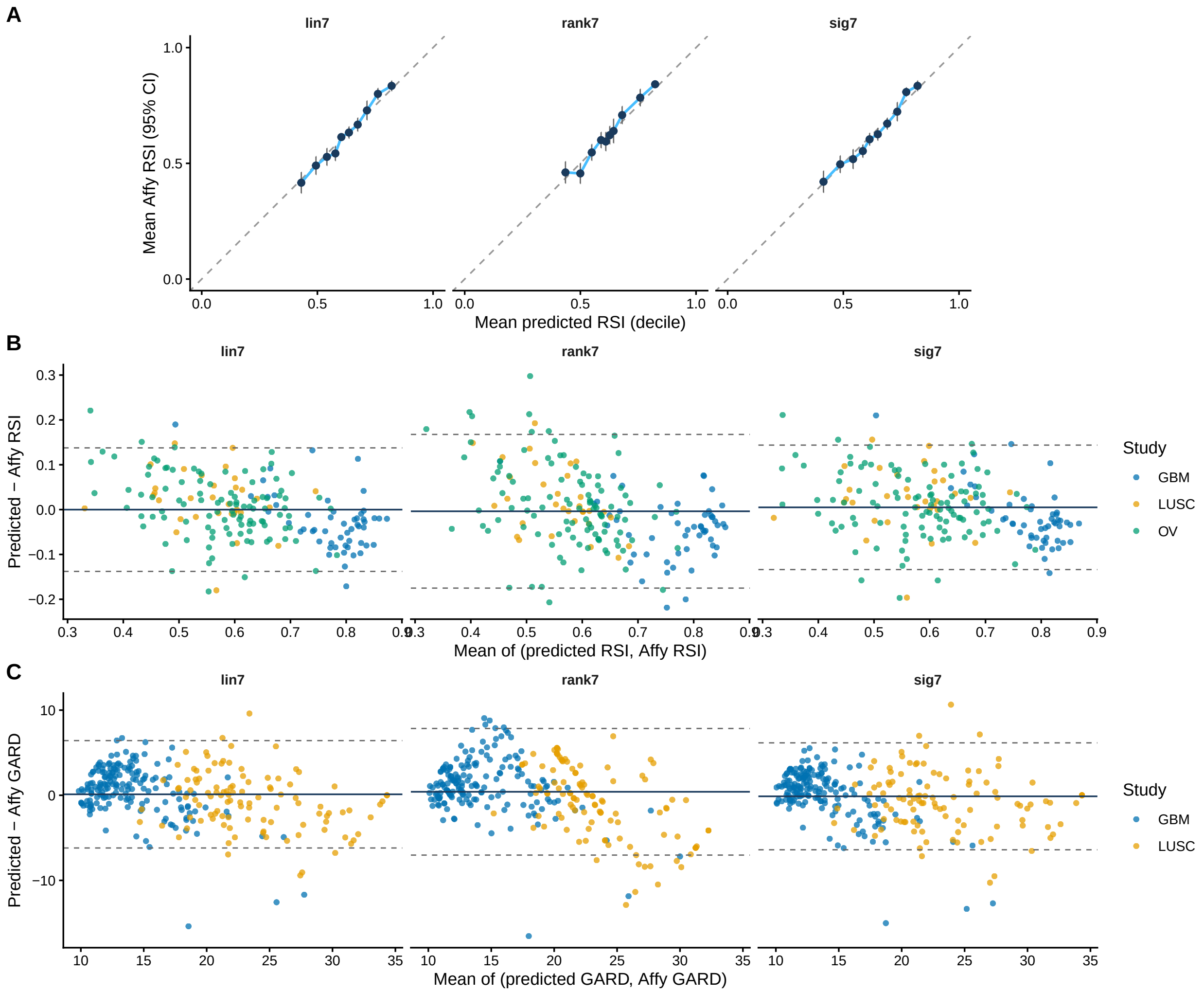


**Figure S3. Variant performance diagnostics.** (**A**) Decile calibration: mean predicted RSI vs. mean microarray RSI per predicted-RSI decile, with 95% CI on the observed mean and the dashed identity line. (**B**) Bland–Altman agreement between predicted and reference RSI: difference (predicted − reference) on the y-axis vs. the mean of the two on the x-axis, with the mean difference (solid) and $\pm 1.96\,\mathrm{SD}$ limits (dashed) per variant. (**C**) Same Bland–Altman construction for predicted GARD vs. GARD-microarray on matched GBM and LUSC samples ($n = 284$).

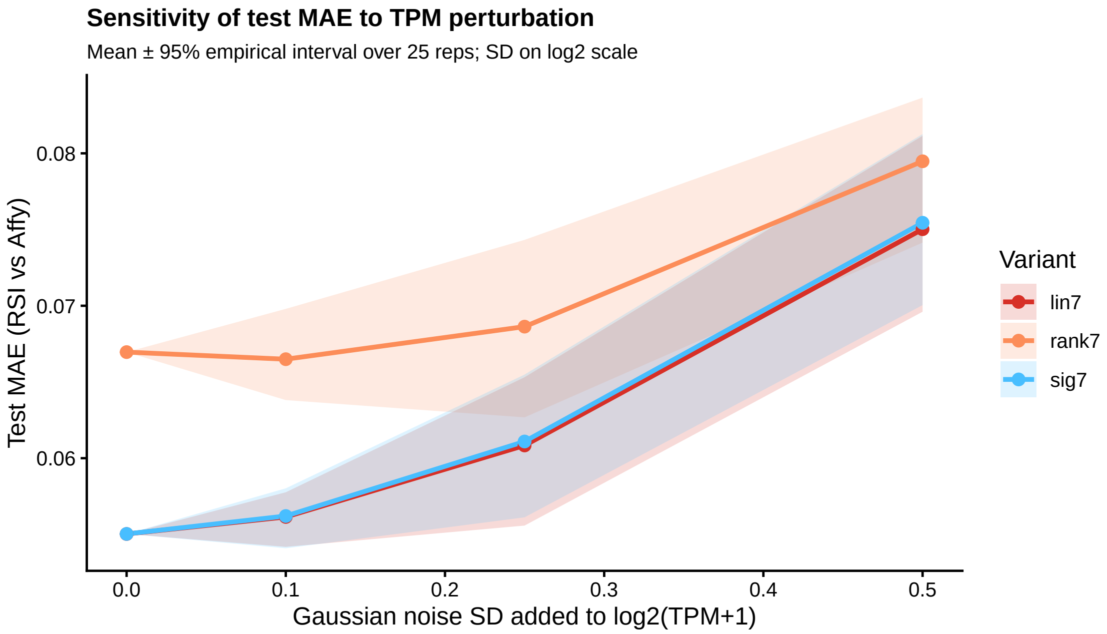


**Figure S4. Sensitivity of test MAE to TPM perturbation.** Test MAE for each variant evaluated under three Gaussian-noise levels added to $\log_2(\mathrm{TPM}+1)$ of the seven modeling genes ($\sigma \in \{0.10, 0.25, 0.50\}$, plus the no-noise reference); solid lines are mean MAE over 25 reps per noise level, ribbons are 95% empirical intervals.

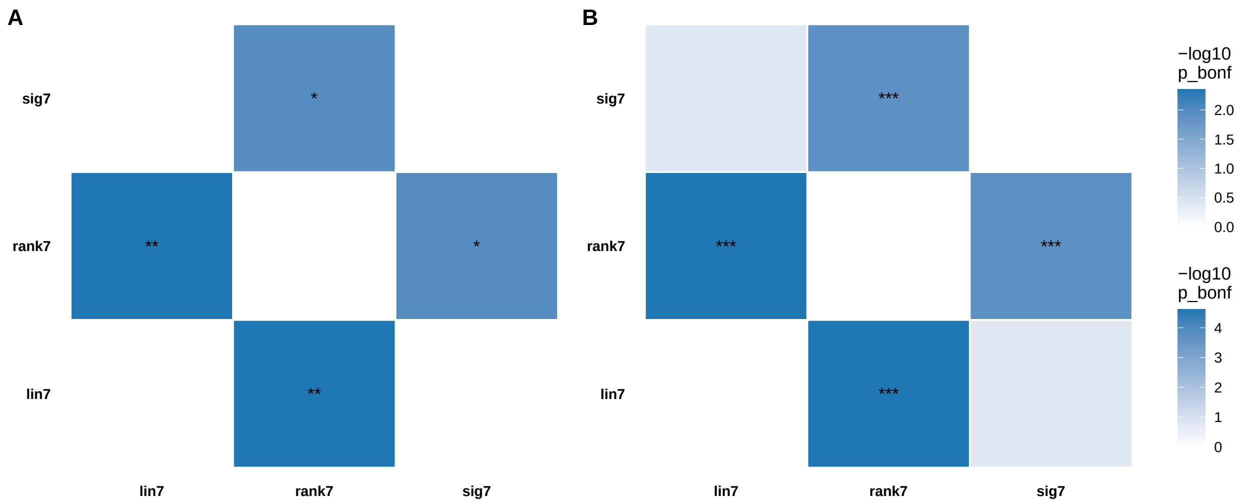


**Figure S5. Pairwise statistical comparison among the candidate variants.** (**A**) Pairwise paired-Wilcoxon tests on $|\text{error}|$ between predicted RSI and microarray RSI; heatmap of $-\log_{10}$ Bonferroni-corrected $p$ values. (**B**) Same construction for Steiger $Z$ tests on the dependent overlapping correlations between each variant's prediction and microarray RSI. * $p < 0.05$, ** $p < 0.01$, *** $p < 0.001$ (Bonferroni-corrected).

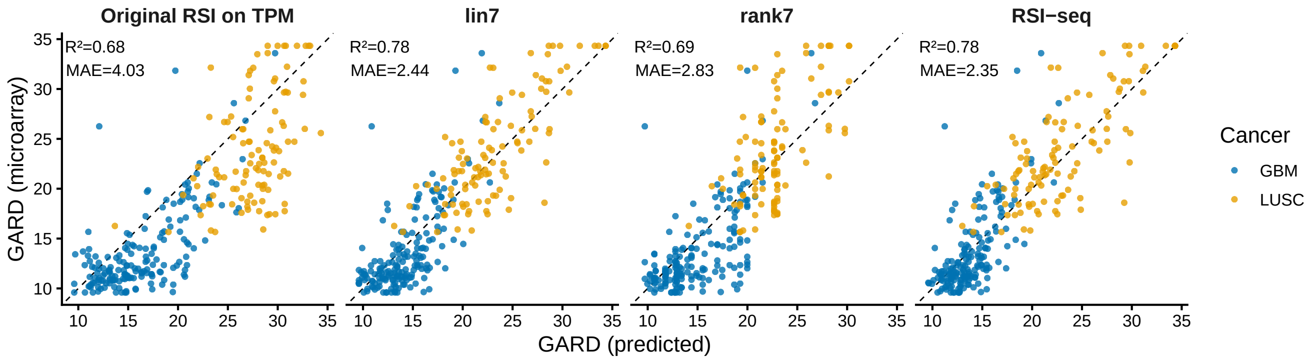


**Figure S6. Per-sample GARD agreement against the microarray reference.** Predicted GARD (60 Gy/30 fx; $\beta = 0.05\,\mathrm{Gy}^{-2}$, $d = 2\,\mathrm{Gy}$) for each candidate model vs. GARD computed from the microarray RSI on the same matched GBM and LUSC samples ($n = 284$); points colored by study.

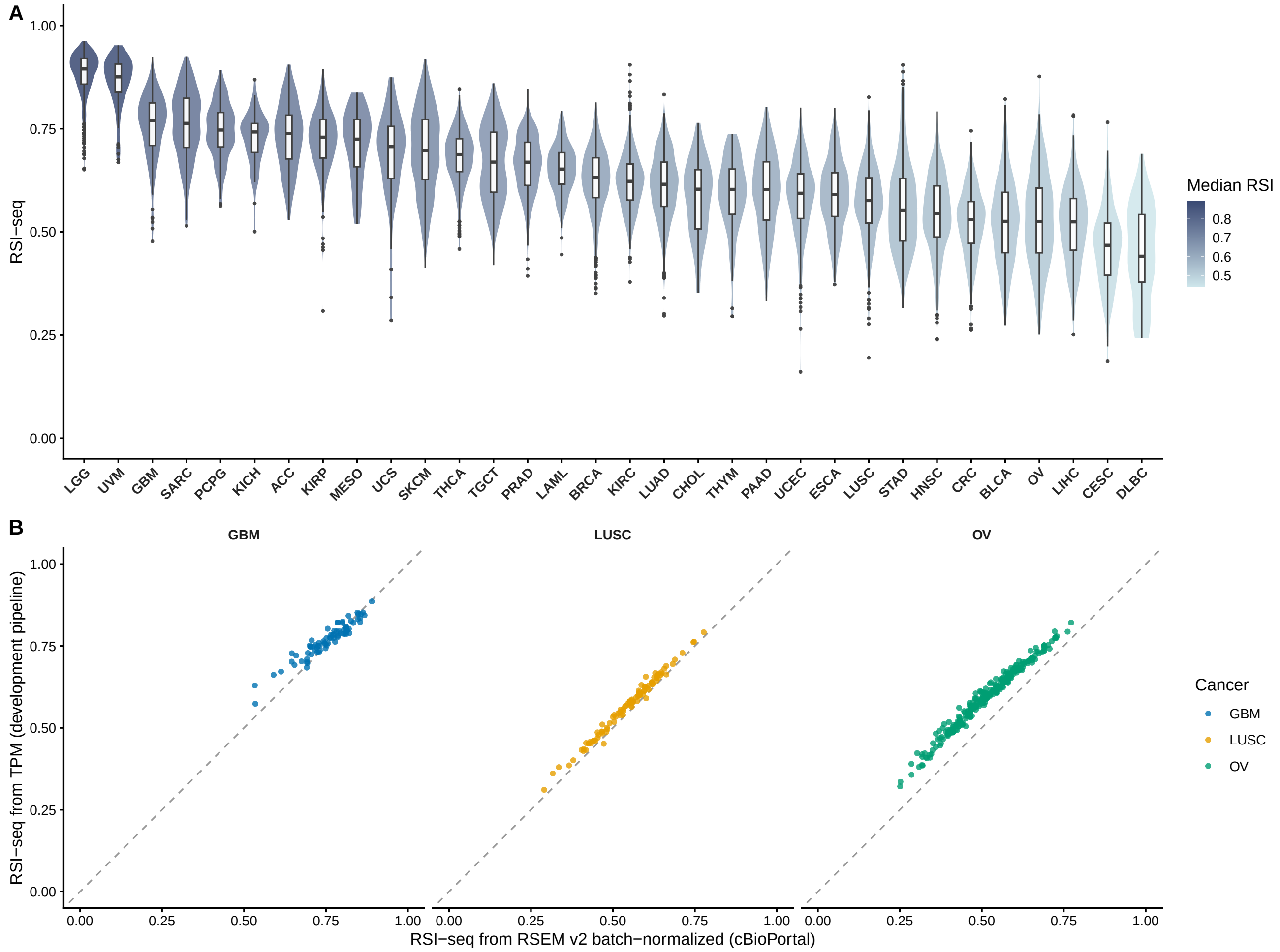


**Figure S7.** **Pan-TCGA application.** (**A**) Per-cancer distribution of RSI-seq computed on the cBioPortal pan-cancer atlas RNA-seq (RSEM v2 batch-normalized; COAD and READ merged into CRC) for all 32 TCGA cohorts. Each cohort is shown as a violin with overlaid boxplot; cohorts are ordered by median RSI-seq from radioresistant (left) to radiosensitive (right), with violin fill encoding the cohort median. The corresponding cohort-level scatter against published microarray RSI medians is shown in **main-text Figure 4**. (**B**) Re-computing RSI-seq on TPM (development pipeline) and on RSEM (cBioPortal) for the same GBM, LUSC, and OV samples gives pooled $n = 437$, Spearman $\rho = 0.97$, Pearson $r = 0.97$, MAE $= 0.056$. Within-sample ranks are well preserved between the two normalizations.

287 ## Supplementary Tables

**Table S1.** Per-study quality-control breakdown.

| Study | $N$ samples | Microarray QC passed | RNA-seq (TPM) QC passed | Both QC passed |
|---|---|---|---|---|
| GBM | 260 | 247 | 179 | 177 |
| LUSC | 130 | 121 | 114 | 107 |
| OV | 417 | 392 | 372 | 353 |

**Table S2.** Cross-platform gene-expression concordance. Sample-wise Spearman correlations between matched Affymetrix (RMA) and RNA-seq profiles, summarized per study. TPM and FPKM produce identical Spearman summaries because Spearman is invariant under monotonic re-normalization; only TPM is reported in the main text.

| Dataset | $n$ samples | $n$ shared genes | Global median $\rho$ | Global SD | RSI-gene median $\rho$ |
|---|---|---|---|---|---|
| GBM (TPM) | 177 | 10,381 | 0.772 | 0.027 | 0.806 |
| GBM (FPKM) | 177 | 10,381 | 0.772 | 0.027 | 0.806 |
| LUSC (TPM) | 107 | 10,381 | 0.803 | 0.015 | 0.806 |
| LUSC (FPKM) | 107 | 10,381 | 0.803 | 0.015 | 0.806 |
| OV (TPM) | 353 | 10,381 | 0.794 | 0.015 | 0.709 |
| OV (FPKM) | 353 | 10,381 | 0.794 | 0.015 | 0.709 |

**Table S3.** RSI model coefficients across formulations. The original 10-gene rank-based RSI is shown in its rank form (Eschrich 2009 coefficients applied to within-sample ranks of the 10 genes). The three RSI-seq candidates use only the seven retained genes; RSI-seq$_{\text{lin}}$ and RSI-seq$_{\text{sig}}$ operate on $\log_2(\text{TPM}+1)$, RSI-seq$_{\text{rank}}$ on within-sample ranks of the seven genes. The final RSI-seq operates on the linear predictor $z$ and maps $\text{RSI} = 1/(1+e^{-z})$. "—" indicates the gene is not in the model.

| Gene | Rank RSI (MA) | RSI-seq$_{\text{lin}}$ | RSI-seq$_{\text{rank}}$ | RSI-seq (final) |
|---|---|---|---|---|
| *AR* | −0.0098 | −0.0029 | +0.0279 | −0.0086 |
| *JUN* | +0.0128 | −0.0018 | +0.0577 | −0.0040 |
| *STAT1* | +0.0255 | +0.0068 | +0.0515 | +0.0426 |
| *PRKCB* | −0.0018 | +0.0121 | +0.0430 | +0.0809 |
| *RELA* | −0.0038 | — | — | — |
| *ABL1* | +0.1070 | +0.0927 | +0.1209 | +0.4211 |
| *SUMO1* | −0.0003 | — | — | — |
| *PAK2* | −0.0092 | — | — | — |
| *HDAC1* | −0.0204 | −0.0517 | +0.0201 | −0.2606 |
| *IRF1* | −0.0442 | −0.0631 | — (NA, dropped) | −0.2924 |
| (Intercept) | — | +0.6779 | −0.7688 | +0.8335 |

RSI-seq$_{\text{lin}}$ and RSI-seq$_{\text{rank}}$ are fit directly against the scaled microarray RSI; the final RSI-seq is fit against $\text{logit}(\text{RSI}_{\text{MA}})$ with $\text{RSI}_{\text{MA}}$ clipped at $\varepsilon = 0.01$ from the boundary. For RSI-seq$_{\text{rank}}$, `lm` dropped *IRF1*'s rank as NA because the seven within-sample ranks always sum to 28 (linear dependence).

**Table S4.** Per-variant test-set performance with bootstrap 95% CIs (1,000 paired non-parametric resamples) on the scaled $[0,1]$ RSI axis. RSI-seq$_{\text{lin}}$ is reported for context but is not deployable (its outputs are not bounded in $(0,1)$, which destabilizes the downstream GARD computation).

| Variant | CV MAE | CV $R^2$ | Test MAE [95% CI] | Test $R^2$ [95% CI] |
|---|---|---|---|---|
| RSI-seq$_{\text{lin}}$ (not deployable) | 0.058 | 0.729 | 0.055 [0.049, 0.061] | 0.78 [0.71, 0.83] |
| RSI-seq$_{\text{rank}}$ | 0.064 | 0.650 | 0.067 [0.060, 0.074] | 0.65 [0.55, 0.73] |
| **RSI-seq (final)** | **0.057** | **0.730** | **0.055 [0.049, 0.060]** | **0.77 [0.69, 0.83]** |

**Table S5.** GARD distribution summary per RSI source and study. 60 Gy in 30 fractions; $\beta = 0.05\,\text{Gy}^{-2}$ and $d = 2\,\text{Gy}$. OV omitted because definitive radiotherapy is not standard in this disease.

| Study | RSI source | $n$ | Mean | Median | SD | Range |
|---|---|---|---|---|---|---|
| GBM | RSI-MA (reference) | 177 | 13.66 | 12.13 | 4.16 | 9.59–33.6 |
| GBM | Original RSI on TPM | 177 | 16.32 | 15.59 | 4.01 | 9.59–29.7 |
| GBM | RSI-seq$_{\text{lin}}$ (not deployable) | 177 | 14.31 | 13.91 | 2.83 | 9.59–23.7 |
| GBM | RSI-seq$_{\text{rank}}$ | 177 | 14.86 | 14.03 | 3.43 | 9.71–26.8 |
| GBM | **RSI-seq (final)** | **177** | **13.93** | **13.45** | **2.44** | 9.59**–**22.7 |
| LUSC | RSI-MA (reference) | 107 | 23.91 | 22.62 | 5.34 | 15.7–34.3 |
| LUSC | Original RSI on TPM | 107 | 27.61 | 28.07 | 3.33 | 13.7–34.3 |
| LUSC | RSI-seq$_{\text{lin}}$ (not deployable) | 107 | 23.21 | 22.53 | 4.41 | 13.1–34.3 |
| LUSC | RSI-seq$_{\text{rank}}$ | 107 | 22.93 | 22.74 | 3.22 | 14.1–30.2 |
| LUSC | **RSI-seq (final)** | **107** | **23.18** | **22.32** | **4.85** | 13.0**–**34.3 |
